# Temporal Dynamics Outweigh Fungicide Treatment in Shaping the Turfgrass Phyllosphere Resistome

**DOI:** 10.64898/2026.06.02.729574

**Authors:** Natalie Wieber, Sara Haney, James Lazarcik, Paul Koch, Jeri Barak

**Affiliations:** Department of Plant Pathology, University of Wisconsin – Madison, Wisconsin, USA; Department of Civil and Environmental Engineering, University of Wisconsin – Madison, Wisconsin, USA

## Abstract

Antibiotic resistance genes (ARGs) are an emerging class of environmental contaminants with significant implications for public health. Previous studies have linked fungicide exposure to elevated levels of ARGs in soil microbiomes, but research investigating the impacts of fungicide on ARGs within phyllosphere bacterial communities is limited. To address this, creeping bentgrass was treated with the fungicide active ingredients chlorothalonil, fluxapyroxad, and propiconazole and sampled at 4 hours, 96 hours, and 168 hours post-application, along with a non-treated control. Quantitative PCR (qPCR) was performed to quantify the abundance of ARGs and a metal resistance gene’s (MRG) abundance relative to 16S rRNA. Additionally, 16S rRNA gene sequencing was performed to characterize bacterial community composition. Results indicated that fungicide treatments did not significantly alter the relative abundance of ARGs or an MRG within bent grass bacterial communities. However, significant changes were observed over time, with changes in ARG and MRG abundance mirroring temporal shifts in bacterial beta diversity. ARGs and the MRG relative abundance had significant correlations with *Proteobacteria, Actinobacteria, Bacteriadota,* and *Firmicutes,* including with genera *Pseudoxanthomas,* and *Dyandobacter,* which contain opportunistic human pathogens. This study demonstrates that fungicides have limited influence on the abundance of ARGs and MRGs in the phyllosphere and helps guide further investigations aiming to mitigate the spread of antibiotic resistance.

**Importance:** Antibiotic resistance makes it harder to treat bacterial infections. Recent evidence suggests that fungicides may increase the abundance of antibiotic resistance genes (ARGs) in soil. Leaves are also treated with fungicides, but it is unclear if there will be increases of antibiotic resistance in this environment because bacteria on leaves face different physiological stresses. Additionally, plants may serve as a route of infection to humans or animals with antibiotic resistant bacteria making it a critical micro-environment to investigate. The purpose of this study was to determine whether the prevalence of antibiotic resistance on plant surfaces changes after short-term exposure to fungicides. This study shows that over short-term application periods, time has a greater effect on the abundance of genes that cause antibiotic resistance than treatment with the fungicides chlorothalonil, fluxapyroxad, and propiconazole on plants. This work helps inform future efforts to mitigate the spread of antibiotic resistance.

## Introduction

Antimicrobial-resistant infections were estimated to result in 1.14 million deaths in 2021 and are projected to increase almost 70% by 2050, with antibiotic resistant bacterial pathogens posing the greatest threat to human health (1). Beyond human health, increased antibiotic resistance may also impact food security by negating available treatments for infected animals and plants (2, 3). Addressing this issue requires a better understanding of the factors that drive antibiotic resistance in various environmental compartments.

Exposure to both clinical and agricultural antibiotics commonly results in higher abundances and increased horizontal gene transfer of antibiotic resistance genes (ARGs) in bacterial communities serve as a significant driver for increases in antibiotic resistance globally (4–6). These increases in ARGs are due to antibiotics selecting bacteria that already carry resistance genes and driving adaptations in bacteria to increase their chances of survival. One example of this selection in nature was shown by Yang et. al. (2021), which found that sulfadiazine, norfloxacin, roxithromycin, and tetracycline concentrations have significant positive correlations to the relative abundance of 25 ARGs in soil (7). Bacteria undergo similar stress responses in the presence of other environmental, chemical, and physical agents, often called non-antibiotic stressors. Non-antibiotic stressors like heavy metals and pesticides have also been shown to increase the abundance and spread of ARG (7–9). This co-selection may be due to shared resistance mechanisms or the activation of shared upstream regulatory pathways. In addition, genes which confer resistance to both types of stressors may be present on a shared mobilizable gene element. In this case, exposure to non-antibiotic stressors may increase horizontal gene transfer through either direct selective pressures or by triggering an SOS response as has been shown previously in soil and *in vitro* (8, 10, 11) For example, Zhu et. al. (2024) found that widely used pesticides cryomazine and kresoxim-methyl led to increased transfer of plasmids carrying ampicillin resistance *in vitro* by increasing membrane permeability as part of an SOS response (8).

Fungicides specifically have been shown to increase the relative abundance of ARGs in soil and worm guts, but it is unclear if they act as drivers for antibiotic resistance in other niches (12–14). Fungicide application is not limited to soil and can impact the bacteria present on plant surfaces (15, 16). The phyllosphere is known to be a harsh environment with limited nutrient availability and intense UV irradiation, leading to lower bacterial density and diversity within this niche when compared to bacterial populations in soil (17–20). Exposure to fungicides may have differing impacts on ARGs in these niches as the diversity and density of bacterial communities alter the spread of ARGs in bacterial communities (21, 22). Huang et al. (2025) previously investigated the impact of applying the fungicides carbendazim and pyraclostrobin to potted lettuce plants in laboratory experiments and found that neither fungicide led to significant shifts in the relative abundance of ARGs throughout the soil-lettuce system (23). Examples of other commonly used fungicides that are relevant to turf include chlorothalonil, propiconazole, and fluxapyroxad. Chlorothalonil is a broad-spectrum fungicide that may act as a bacteriostatic due to its ability to inhibit glutathione (24). Additionally, chlorothalonil has been shown to disrupt bacterial communities (16, 25, 26), and enhance the abundance of some ARGs in soil and in earthworms (12, 13). Fluxapyroxad and propiconazole act by inhibiting succinate dehydrogenase and ergosterol synthesis, respectively, and do not exhibit strong antibacterial potential. To our knowledge neither fluxapyroxad nor propiconazole have been previously tested for impacts on ARG abundance.

Amenity turfgrass found on golf courses is an especially relevant host for this study as it is intensively managed and heavily treated with fungicides to prevent disease (27–29). Models estimate that the most concentrated fractional areas of turf grasses align with major cities (30). Observing ARG abundances in this environment is important since bacteria containing ARGs are not confined to their plant host and contact with urban green spaces increases microbial diversity of human microbiomes via introduction of bacteria from the environment (31, 32) This finding suggests that humans interacting with intensively managed turfgrass landscapes could have higher exposures to ARGs, though the risk for increased antibiotic resistant illnesses as a result of these exposures remains uncertain.

This field study tested the hypothesis that acute fungicide treatments have off-target impacts on the phyllosphere bacterial community. Changes to the relative abundance of ARGs and the bacterial community composition were measured using HT-qPCR and 16s rRNA sequencing. In addition to abundance and composition, we tested the hypothesis that fungicide treatments can drive off-target changes to ARG associations within the bacterial community.

## Methods

### Field design, treatment, and sampling

This study was carried out at the OJ Noer Turfgrass Research Facility in Madison, WI, USA during the summer of 2022 on creeping bentgrass maintained at a length of 1.27 cm. The study included a non-treated control and six treatments comprised of commercial formulations of chlorothalonil, fluxapyroxad, and propiconazole fungicides applied alone or in combination with each other. Plots measured 0.9 by 1.5 meters and were arranged within a randomized complete block design with four replications. Treatments were applied at a nozzle pressure of 276 kPa using a CO2-pressurized sprayer equipped with one Teejet AI9508EVS nozzle. All fungicides were agitated by hand and applied at the equivalent of 5.7 L of water per 92.9 m^2^. Treatment dosage and reapplication frequency were based on the label recommendations for each product and can be found in Table 1. Treatment applications began on June 1^st^, 2022, and sampling began on July 5^th^, 2022. The 0 hour, 4 hour, 96 hour, and 168h timepoints mentioned in this paper are relative to the application that occurred on July 5^th^, 2022.

**Table 1.**
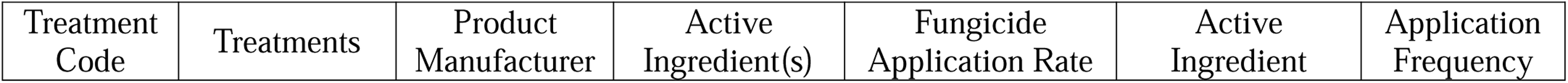
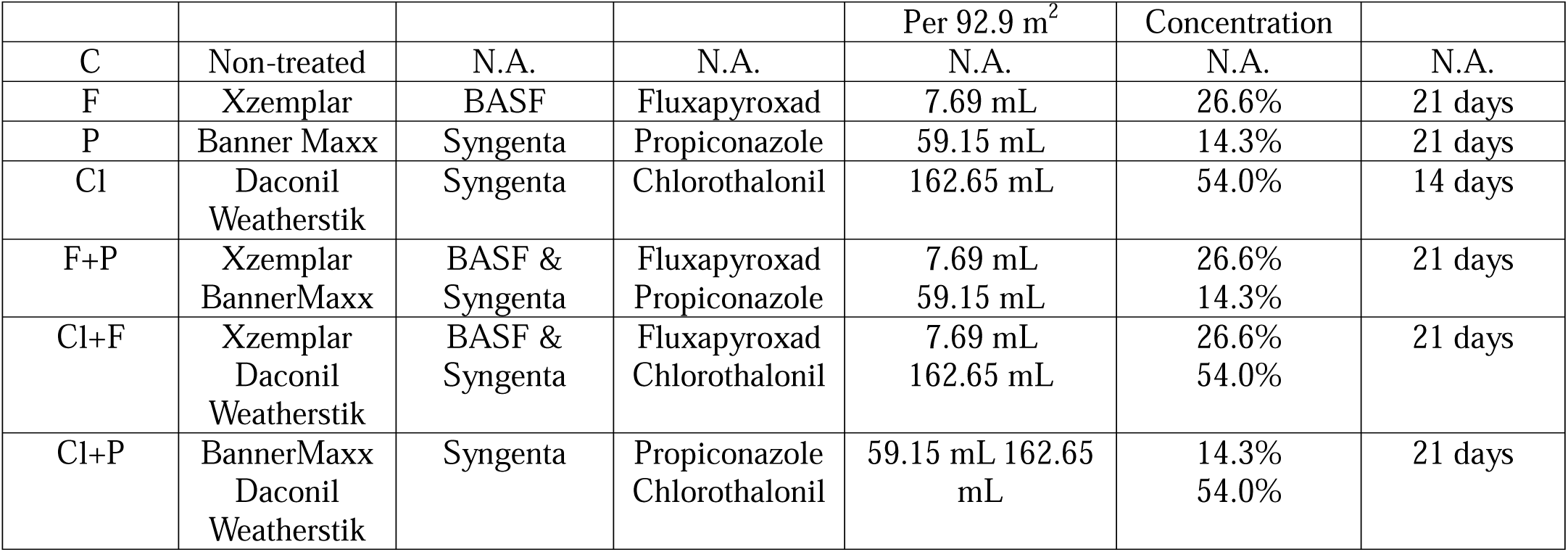
Field Treatments, Active Ingredients, and Application Rates.

Turf clippings were collected from each plot prior to treatment with fungicide for the 0 hour (0h) sample and then at 4 hours (4h), 24 hours (24h), 96 hours (96h), and 1 week (168h) post application. To collect clippings, a mower set to 0.2 mm was driven over the plot. Clippings were collected from a basket attached to the mower and transferred into plastic bags. Between each plot, gloves were sprayed with 70% ethanol and the basket was brushed free of remaining turf clippings. Samples were put into sealed plastic bags and placed on ice for less than 1.5 hours during the completion of sampling and transport back to the laboratory. Samples were then stored at -80°C until further processing.

### DNA extraction

Phyllosphere bacterial DNA was extracted from 5 g of turf clippings for each sample using a benzyl chloride liquid: liquid extraction followed by an isopropanol precipitation, as described previously by Suda, Bowsher, and Grandy et al. (33–35). Briefly, leaf clippings were combined with 5 mL extraction buffer (100 mM Tris-HCl, pH 9.0, 40 mM EDTA), 1 mL 10% SDS (Life Technologies, Carlsbad, CA) and 3 mL Benzyl Chloride (Sigma Aldrich, St. Louis, MO). This was then incubated at 50°C for 15 minutes with each sample being briefly vortexed at 1-minute intervals. Leaf tissue was then removed from the solution using tweezers and a fine mesh sieve that were cleaned with 70% ethanol between samples. The solution was then mixed with 3 mL 3M sodium acetate (Sigma Aldrich, St. Louis, MO), vortexed, incubated on ice for 10 minutes, then centrifuged at 6000 g for 15 minutes at 4°C. The aqueous layer was moved into a new 50 mL conical tube and centrifuged at 6000 g for 15 minutes at 4°C. The aqueous layer was again moved into a new 50 mL conical tube, mixed with an equal volume of isopropyl alcohol, inverted several times to precipitate the DNA, and then centrifuged at 6000 g for 15 minutes at 4°C. The supernatant was discarded, the pellet was allowed to air dry for 20 minutes and was stored in ethanol (Decon Labs, King of Prussia, PA) overnight at -20°C. The pellet was resuspended in ethanol and centrifuged at 16,000 g for 15 minutes. The supernatant was removed and the pellet was resuspended in TE buffer (10mM Tris-HCL 1mM EDTA). DNA was stored at -80°C for qPCR and 16S rRNA sequencing.

### SmartChip Analysis

DNA was diluted to 10 ng/uL using sterile water and sent for high-throughput qPCR at the Resistomap Oy Laboratory (Resistomap Oy, Helsinki, Finland) using the SmartChip Real Time PCR system (Takara Bio, CA, USA). Samples were analyzed using a customized primer set targeting seven genes, see Table 2 for primer sequences. Primers were initially selected based on a preliminary screen of four field samples using 384 available primer sets targeting ARGs, mobile genetic elements, and metal resistance genes (MRGs), which were narrowed down to 24 genes of interest (36). Genes of interest were chosen for further screening based on a greater than 10% difference in gene abundance between treatment and time groups, and because they confer resistance to antibiotic classes listed in the 2017 WHO Global Priority Pathogen List (37). Of the 24 genes screened only seven are included in this paper. Those not included often fell below the limit of detection leading to sparse, non-parametric data sets. While *czcA* is a metal resistance gene (MRG), its ability to confer resistance to multiple metals, positive associations to multiple antibiotic resistance classes, and results during the preliminary screening led to its inclusion in this study (38–40).

**Table 2.**
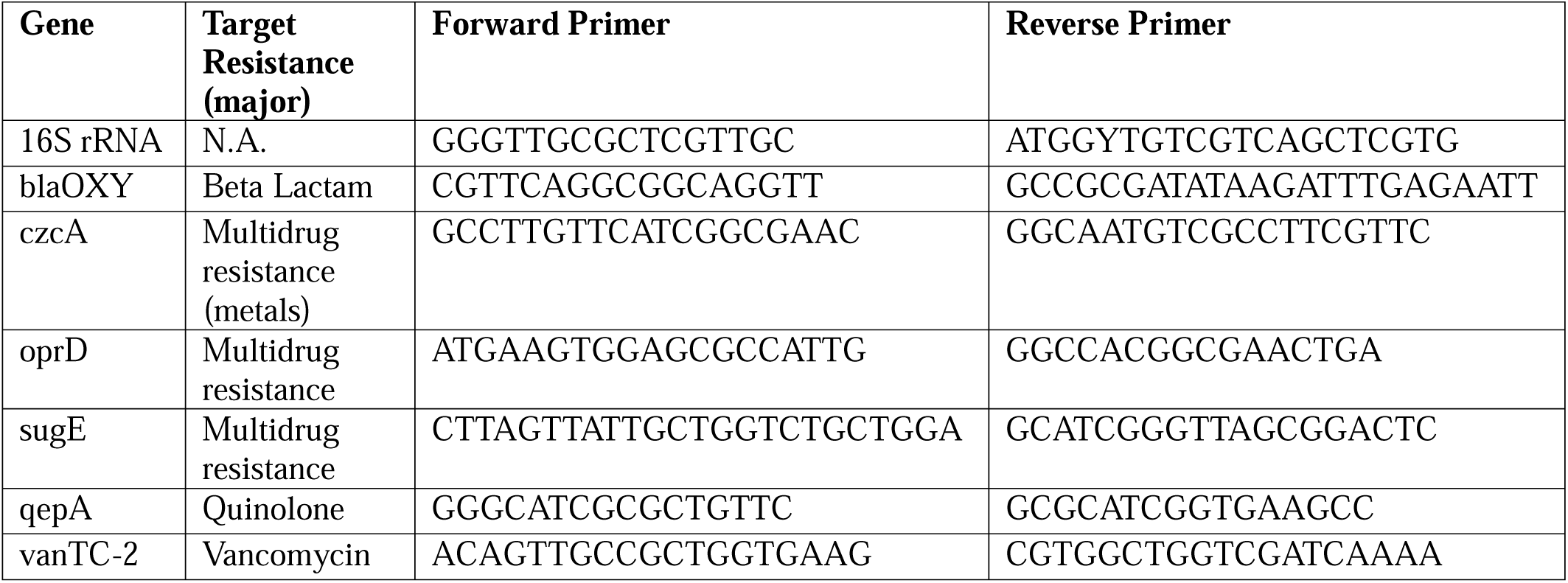
Primer Targets and Sequences (36) for SmartChip HT-qPCR Analysis.

### 16S rRNA amplification and sequencing

The variable 4 (V4) region of the 16S rRNA gene was sequenced using one-step barcoded primers (41). DNA was diluted to 25-5 ng/ µL using sterile water. To amplify the V4 region, 1 µL template DNA was combined with 6.25 uL, 0.5uM of Platinum II Hot Start Master Mix (2X) (Thermo Fisher, Waltham, MA), 0.6 µL of 5 um mitochondrial and chloroplast clamps (42), 1 µL of 3% BSA (Sigma Aldrich, St. Louis, MO), 2.5 µL platinum II GC enhancer (Thermo Fisher, Waltham, MA), sterilized water was added to increase volume to 25 µL. PCR cycling conditions initially held samples at 94°C for 3 minutes and then had 30 cycles with 15 seconds at 94°C, 10 seconds at 75°C, 10 seconds at 55°C, 1 minute at 68°C and 10 seconds at 72°C. Amplicons were run on a 1.5% agarose gel to check that they were the expected size and then gel extracted with the Zymoclean Gel DNA recovery kit (Zymo Research, Irvine, CA). Gel extracted DNA was quantified with the Qubit dsDNA HS assay (Thermo Fisher, Waltham, MA) and pooled into an equimolar 4 nM solution prior to sequencing on an Illumina MiSeq (llumina, San Diego, CA, USA) system with a 2×250 kit with a 10% PhiX control (llumina, San Diego, CA, USA).

### Fungicide solid phase extraction

Propiconazole and fluxapyroxad were extracted from turf samples using methods previously established by González et al. (2022) and Chen et al. (2016), respectively (43, 44). Two hundred milligrams of each frozen turf sample was weighed into an individual MP biomedical lysing matrix D tube (MP Biomedicals, Irvine, CA). Propiconazole-D5 in acetonitrile (Dr. Ehrenstorfer, Augsburg, Germany) was added as an internal standard at a volume that would equate to a concentration of 5 or 1 ng/mL post extraction and dilution. The propiconazole-D5 was left to adhere to the leaf tissue for five minutes and then 80:20 ACN:MilliQ H_2_O (v/v) was added to the tube for a total volume of 1 mL. Samples were placed in an MP Biomedicals FastPrep-24 benchtop homogenizer and macerated at 6 m/s for 40s. Excess water was removed from the samples by adding 50 mg magnesium sulfate (Chem-Impex International, Wood Dale, IL) and 15 mg of anhydrous sodium acetate (Chem-Impex International, Wood Dale, IL). Samples were then macerated again using the MP Biomedicals FastPrep-24 benchtop homogenizer method. Samples were centrifuged at 5000 g, the liquid phase was transferred into a 2 mL d-SPE tube (Agilent, Santa Clara, CA) and vortexed at 2000 rpm for one minute and centrifuged a third time at 5000 g for 5 minutes. The liquid phase was then transferred into a 2 mL amber vial (Agilent, Santa Clara, CA) for LC-MS/MS analysis. Samples were diluted with 80:20 ACN:H2O (v/v) as needed to fall within the standard curve.

Chlorothalonil was extracted from turf samples as described previously by Hockemeyer et al. (2024), with minor alterations (45). Briefly, 0.2 g of each frozen turf sample was weighed into an individual MP Biomedical lysing matrix D tube (MP Biomedicals, Irvine, CA). Twenty-five microliters of sulfuric acid (Fisher Scientific, Hampton, NH) were added directly to the turf clippings, and nicarbazin (Sigma Aldrich, St. Louis, MO) in acetonitrile (Fisher Scientific, Hampton, NH) was added as an internal standard at a volume that would equate 1 ng/mL post extraction and dilution. The nicarbazin was left to adhere to the leaf tissue for five minutes and then 80:20 ACN:MilliQ H_2_O (v/v) was added to the tube to a total volume of 1 mL. Samples were placed in a MP Biomedicals FastPrep-24 benchtop homogenizer and macerated at 6 m/s for 40 s. Excess water was removed from the samples by adding 50 mg magnesium sulfate (Chem-Impex International, Wood Dale, IL) and 15 mg of anhydrous sodium acetate (Chem-Impex International, Wood Dale, IL). Samples were then macerated again using the same MP Biomedicals FastPrep-24 benchtop homogenizer method above. Samples were centrifuged at 5000 g and the liquid phase was transferred into a 2 mL amber vial (Agilent, Santa Clara, CA) for LC-MS/MS analysis. Samples were diluted with 80:20 ACN:H2O (v/v) as needed to fall within the standard curve.

### LC-MS/MS Methods

Propiconazole was analyzed as previously described by González et al. (2022) using an Agilent 1260 HPLC coupled to a 6460 tandem quadrupole mass spectrometer (43). Propiconazole reference materials were prepared at concentrations of 5, 10, 25, 50, and 100 ng/mL to serve as a standard curve. A single-point propiconazole-d5 reference was included with each run for downstream recovery analysis. An isocratic elution method was performed using an Agilent Poroshell 120 EC-C18 column (3.0 mm x 50 mm, 2.7 µm particle size) and 40% solvent A (0.1% (v/v) formic acid 10% (v/v) acetonitrile in MQ water) and 60% solvent B (100% acetonitrile) at a constant flow rate of 0.25 mL/min. The column temperature was set to 30°C. With an injection volume of 5 µL, propiconazole and propiconazole-d5 were measured with positive polarity after the sample was ionized with an electrospray ionization ESI source, Relevant transitions for propiconazole and propiconazole-d5 are described in Table 3.

**Table 3:** Transition Information for Propiconazole and Propiconazole-d5 LC-MS/MS Method.

| *Compound Name* | *Precursor (m/z)* | *Product (m/z)* | *CE (V)* |
| --- | --- | --- | --- |
| *Propiconazole* | 342.1 | 158.9 | 40 |
|  | 342.1 | 123 | 70 |
|  | 342.1 | 69.1 | 20 |
|  | 342.1 | 41.1 | 44 |
| *Propiconazole-d<sub>5</sub>* | 347.1 | 159 | 36 |
|  | 347.1 | 74.1 | 24 |
|  | 347.1 | 44.1 | 56 |
|  | 347.1 | 43.1 | 44 |

Fluxapyroxad was quantitatively analyzed as previously published by Chen et. al, (2016) but performed on an Agilent 1290 Infinity II coupled to a tandem quadrupole mass spectrometer (44). Fluxapyroxad reference materials were prepared at concentrations of 5, 10, 25, 50, and 100 ng/mL to serve as a standard curve. A single-point propiconazole-d5 reference was included with each run for downstream recovery analysis. Separation was performed using an Agilent Poroshell 120 EC-C18 (3.0 mm x 50 mm, 2.7 µm particle size) column set to 40°C at a constant flow rate of 0.3 mL/min with an injection volume of 5 µL. Solvent A (0.1% formic acid in Milli-Q water) and solvent B (100% acetonitrile) were used as a gradient program for separation. Fluxapyroxad was quantified using a negative ESI while propiconazole-d5 was analyzed with positive ESI at the transitions found in Table 4.

**Table 4:** Transition Information for Fluxapyroxad, and Propiconazole-d5 LC-MS/MS Method.

| *Compound Name* | *Precursor (m/z)* | *Product (m/z)* | *CE (V)* |
| --- | --- | --- | --- |
| *Fluxapyroxad* | 380.1 | 247.9 | 20 |
|  | 380.1 | 130.8 | 20 |
|  | 380.1 | 90.9 | 43 |
| *Propiconazole-d5* | 347.1 | 159 | 36 |
|  | 347.1 | 74.1 | 24 |

### LC-MS/MS method performance

Recovery experiments were performed to assess the accuracy of each method. Four replicates of turf clippings were spiked with either chlorothalonil, propiconazole, or fluxapyroxad, and the corresponding internal standard and extracted as described above. Extraction efficiencies averages were calculated to be 125%, 87%, and 85% for native chlorothalonil, propiconazole, and fluxapyroxad respectively. Nicarbazin internal standard recovery across all chlorothalonil runs was 96% on average (Fig. S1). Propiconazole-D5 average internal standard recovery was 118% and 121% for the propiconazole and fluxapyroxad runs respectively (Fig S1). Standard range linearity for all runs was determined to be R^2^> 0.990 for each compound.

## Data analysis

Fungicide concentrations were calculated as described previously by González et. al (2022) (43). LC-MS/MS signals were corrected based on internal standard recovery, and micrograms of active ingredient per gram of turf were calculated. Samples were averaged across triplicate extractions for reporting, some samples were averaged over only two replicates due to limited turf material. Paired and unpaired Welch’s t-tests were applied to determine significant differences across time and treatment groups respectively.

HT-qPCR samples with 16S rRNA Ct values greater than 18 were removed from the data set over concerns of low overall bacterial DNA concentrations. Ct values greater than 26 were filtered out of the data set due to increased likelihood of false positive signals (36). Additionally, any ARG primer result that had a primer efficiency with a greater than 10% difference from the 16S rRNA efficiency was removed from the data set. Relative abundance was calculated with respect to 16S rRNA using the delta Ct method (46). Changes in ARG abundance were analyzed using linear mixed effect models to account for repeated measurements of the same plots over time. Models were made in R (4.3.2) using the lme4 package (47). Linear mixed effect models fit with ARG abundance as the outcome, and time, treatment, and their interaction as fixed predictors. A random effect of plot was added to account for the correlation of observations on the same plot over time. Residuals were visually inspected for normality and homoscedasticity. Post hoc comparisons were performed using the emmeans package with a Tukey correction. Visualizations were created using ggplot2 (48).

Sequencing reads were quality filtered and cleaned using the DADA2 pipeline (1.32.0) to generate amplicon sequence variants (ASVs) in R (4.3.2). Default parameters were used apart from truncation lengths, which were set to (245,150), and the use of pseudo pooling. Taxonomic ranks were assigned with SILVA (v138) (49, 50). Sequences were normalized using the CSS method in R (4.3.2) (51). Bray Curtis Dissimilarity, Chao diversity, Shannon diversity and permutate-multivariate analysis of variance (PERMANOVA) were performed with the “vegan” and “phyloseq” packages in R (4.3.2) using versions (2.7-1) and (1.46.0) respectively (52, 53). Pairwise comparisons were performed using pairwise adonis (54).

Correlations between treatment, ARGs, MRGs, and phyla were investigated using redundancy analysis using the “vegan” package (2.7-1) in R (4.3.2) (52, 53). Treatment and phyla served as exploratory values, ARGs and the MRG were the response variables. Bidirectional stepwise selection was performed on the initial model containing all variables using the step function from base R (4.3.2). Model terms were iteratively added or removed using RDA-specific constrained ordination methods from the “vegan” package (2.7-1) (52, 53). Candidate models were compared using permutation tests. This process led to the inclusion of *Proteobacteria, Latescibacterota, Gemmatimondota, Methylomirabilota, Chloroflexi*, *Verrucomicrobiota, NB1. J, Acidobacteriota, Plantomycetota, Adidtobacteriota, Cyanobacteria, Armatimonadota, Sumerlaeota, Actinobacteriota,* and *Firmicutes,* as exploratory variables within the final model. Variables were tested for co-linearity through variance inflation factors and significance was determined with permutation tests within the “vegan” package (2.7-1) (53). Spearmen correlations were calculated in base R to assess pairwise correlations between ARGs, MRGs, and genera. Correlations were calculated between ARGs but not between genera to conserve statistical power. Bonferroni corrections were applied to p values. Significant spearman correlations were visualized in Gephi (0.10.1) (55). In all cases statistical significance was defined as p<0.05.

## Results

### Time, not treatment, impacts the resistome and bacterial community within the phyllosphere

Exposure to fungicide had no significant impact on the relative abundance of individual ARGs or MRGs within turf phyllosphere bacterial communities (0.83<P > 0.08) (Fig.1). Therefore, treatment groups were combined for subsequent analysis. Time did have a significant (p < 0.05) effect on the relative abundance of each measured gene (Fig. 2), when referred to as a group these genes will called the phyllosphere resistome moving forward. The overall relative abundance of ARGs and MRGs followed a temporal trend where there was a significant difference between the 0 hour and 4 hour timepoints from those at 168 hours post fungicide application. These results suggest that while the phyllosphere resistome is not impacted by treatment with chlorothalonil, fluxapyroxad, or propiconazole-based fungicide within our field study another variable causes a significant decrease in each of the ARGs and MRGs from our pre-treatment (0h) timepoint and 168 hours post fungicide application.

**Figure 1.**
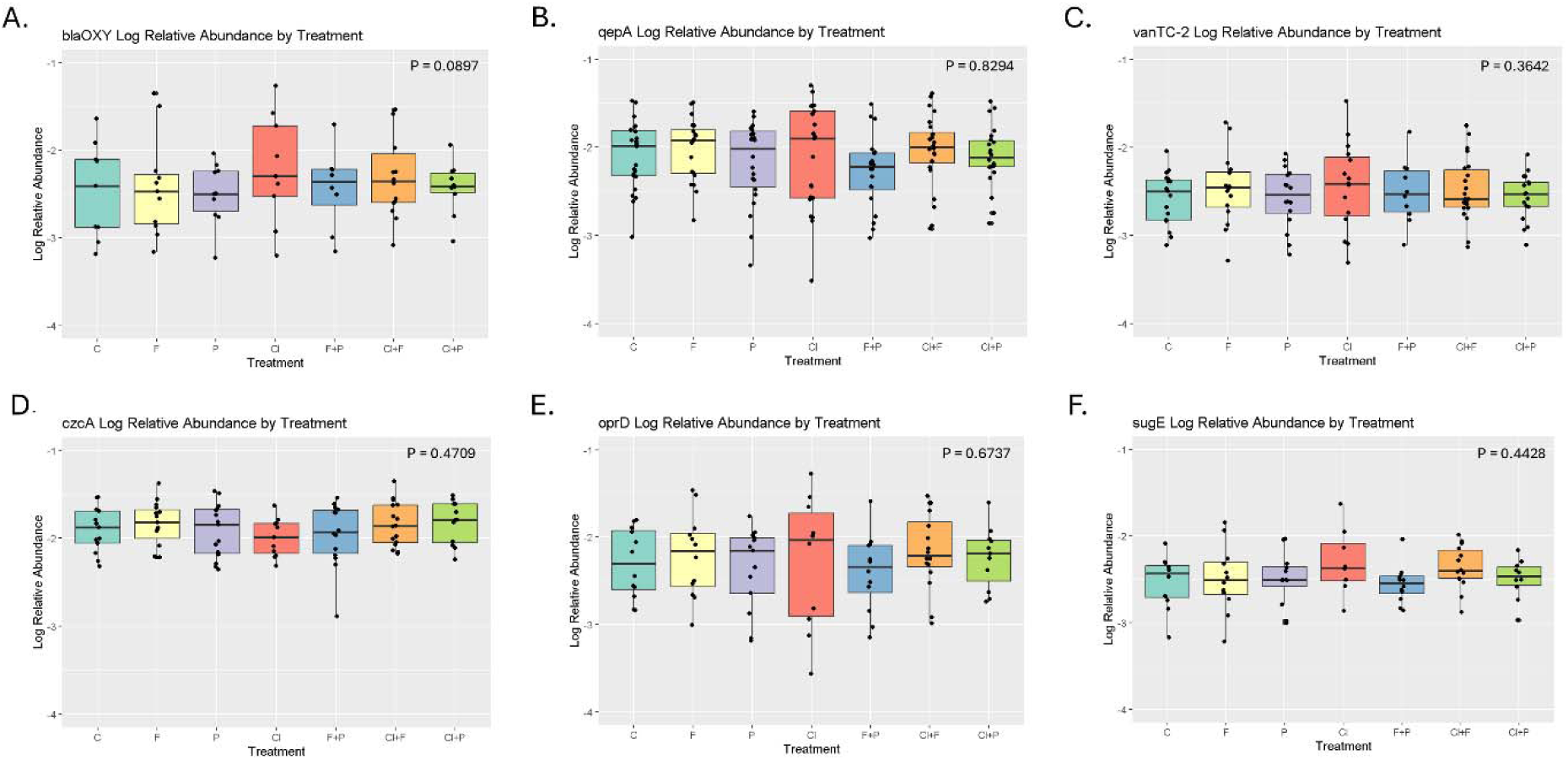
Log relative ARG abundances in the control plot (C) and plots treated with fungicides containing Propiconazole (P), Fluxapyroxad (F), and Chlorothalonil (Cl) as singular treatments or in combination averaged across time points for A. blaOXY, B. qepA, C. vanTC-2, D. czcA, E. oprD, F. sugE.

**Figure 2.**
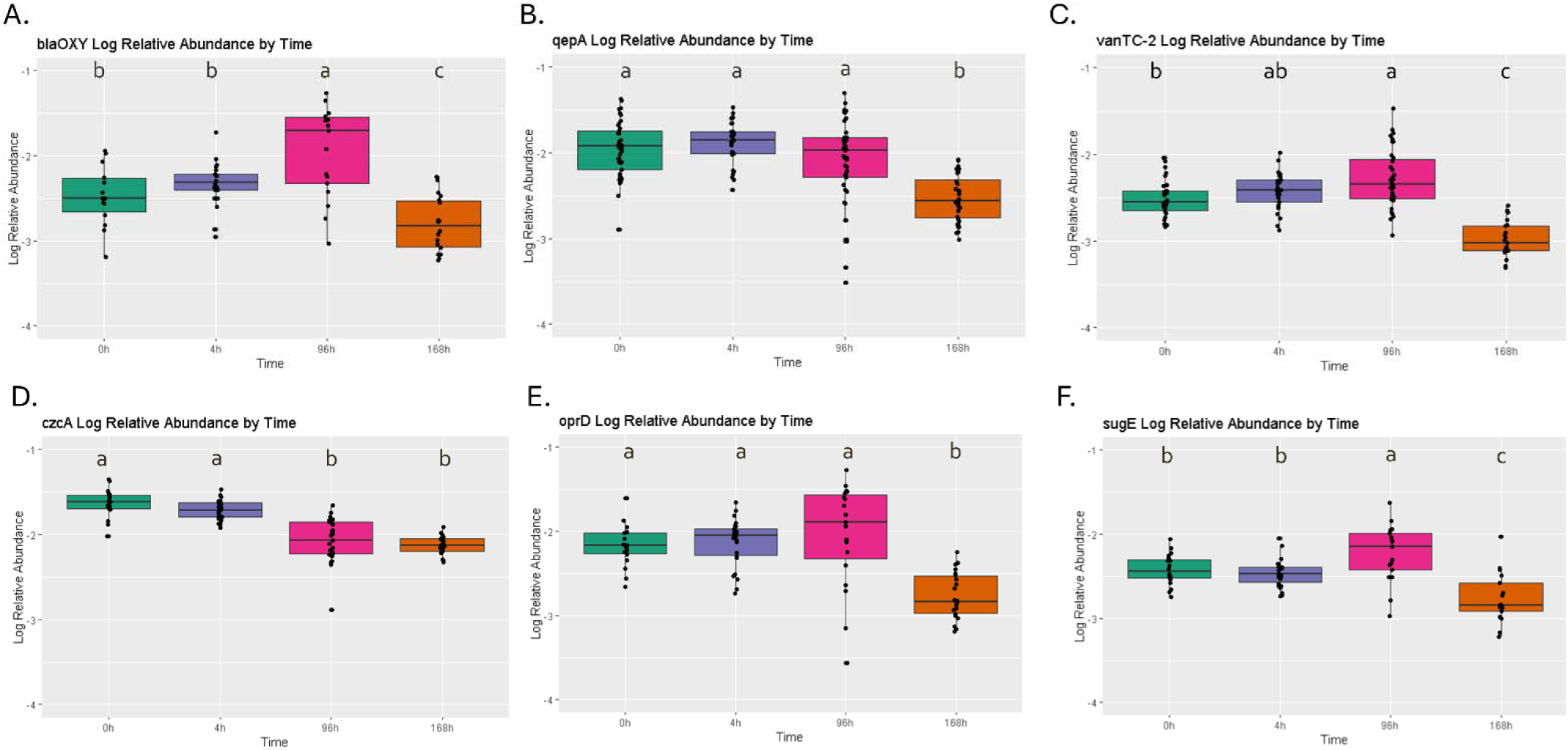
Log relative abundance of ARGs and MRGs across sampling timepoints at 0 hours (0h), 4 hours (4h), 96 hours (96h) and 1 week (168h) post fungicide application for A. blaOXY B. qepA C. vanTC-2 D. czcA E. oprD F. sugE. Abundance is relative to 16S rRNA and was calculated using the delta ct method. Each data point is the average of three technical replicates. Letters denote significance groups within each gene (p<0.05) based on a linear mixed effects Anova followed with a Tukey correction.

After establishing that fungicide treatment did not alter the resistome of the turf phyllosphere, we considered factors that may help characterize the significant change in ARGs and MRGs over time, including changes in bacterial community composition. To do this, we performed sequencing targeting the V4 region of 16S rRNA. A total of 4,952,096 raw reads were generated during sequencing for 107 samples, averaging 46,281 reads per sample. After quality filtering using the DADA2 pipeline, 2,083,477 reads were retained among the remaining 97 samples. In this study, a total of 2,837 ASVs were classified into 29 unique phyla. Rarefication curves of ASVs were saturated for all samples (Fig. S2A) and many samples had good’s coverage values greater than 90% (Fig. S2B), suggesting that sequencing depth appropriately represented the bacterial communities present. After CSS normalization the minimum good’s coverage score was 97.88%.

Bray-Curtis distances were calculated based on ASVs and plotted on multivariate ordination axis (Fig. 3). Fungicide treatments, combined across timepoints, had no significant impact on community structure according to PERMANOVA tests (p > 0.05; Fig. 3A), so treatment groups were combined for subsequent analysis. Time had a significant effect on the beta diversity of samples, accounting for 7.3% of the overall variation according to PERMANOVA analysis, thus leading to varied clustering (Fig. 3B). Pairwise PERMANOVA analysis revealed the communities at the 0 hour and 4 hour timepoints differ from those at 1 week post fungicide application significantly, showing that bacterial community composition is significantly changed over time. The results show that the overall composition of the communities is resilient to our treatment but does change over time. Additionally, we see that the temporal significance mirrors the temporal trend seen in the abundance of ARGs and MRGs where 0 hour and 4 hour time points were significantly different from the 168 hour post fungicide application timepoint. The relative abundance of resistance genes at 96 hours does not follow a shared trend. However, we do see the most variance at 96 hours (Fig. 2, Fig. 3). The increased variability in the resistance gene relative abundances and their deviation from a shared temporal trend may be reflective of the increased community variation among samples within this timepoint. Shared temporal trends across the bacterial community composition and the resistome suggest that they are being influenced by similar factors or are influencing each other.

**Figure 3.**
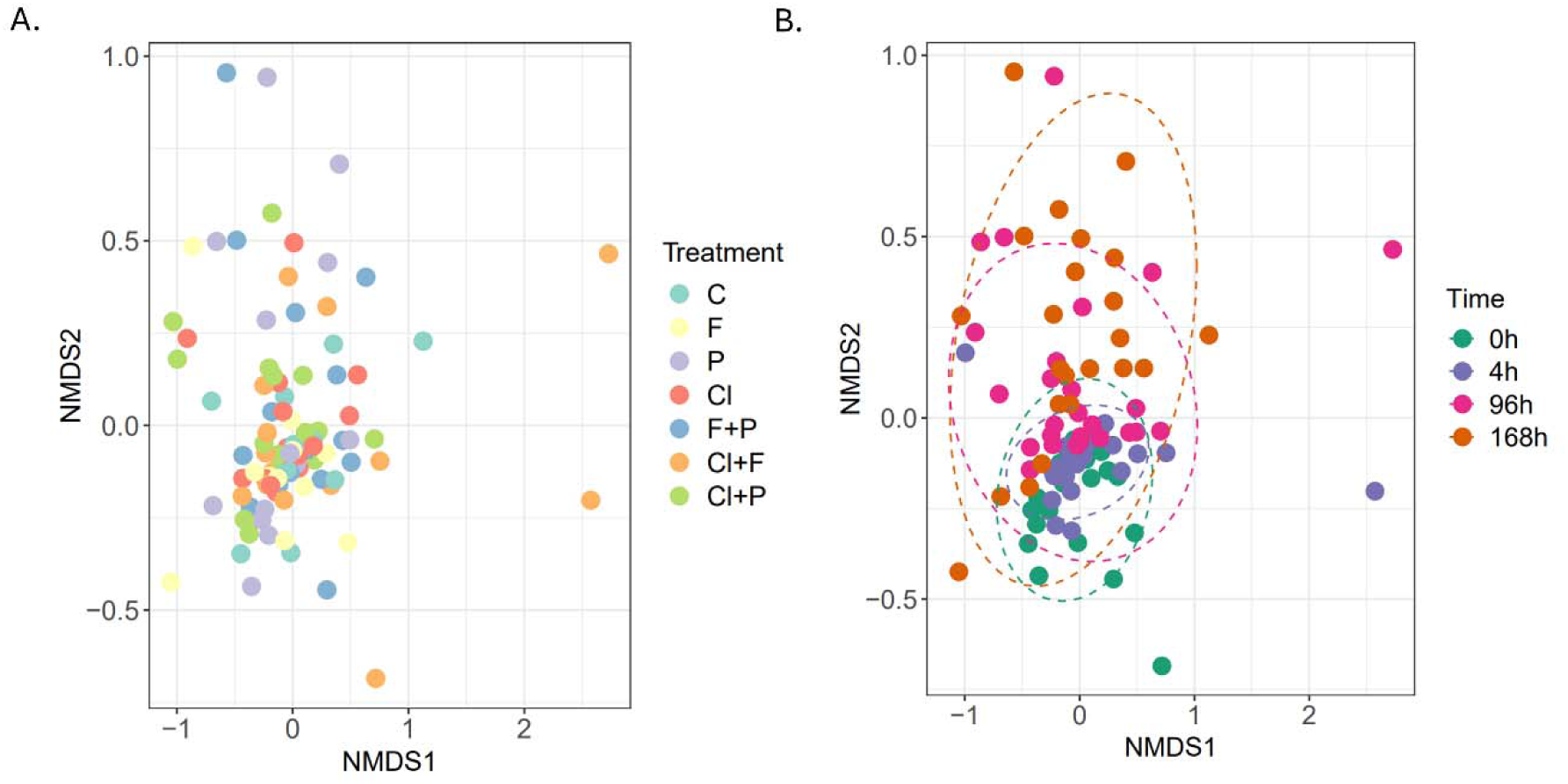
Non-metric multidimensional scaling (NMDS) ordination plot of Bray-Curtis dissimilarities demonstrating the composition of microbial communities across variables. Points represent individual turf samples. A. Treatment groups account for 5.8% of the variability in beta diversity among samples and show no statistical difference between untreated plots (C), and plots treated with Propiconazole (P), Fluxapyroxad (F), and Chlorothalonil (Cl) as singular treatments or in combination. B. Timepoints account for 7.3% variability at 0 hours (0h), 4 hours (4h), 96 hours (96h), and 168 hours(168h) post fungicide application. There is a significant difference between the 168 hourtimepoint when compared with the 0 hour and 4 hour timepoints as determined by a PERMANOVA test and pairwise adonis comparisons. Ellipses represent 95% confidence intervals for group centroids.

**Figure 4.**
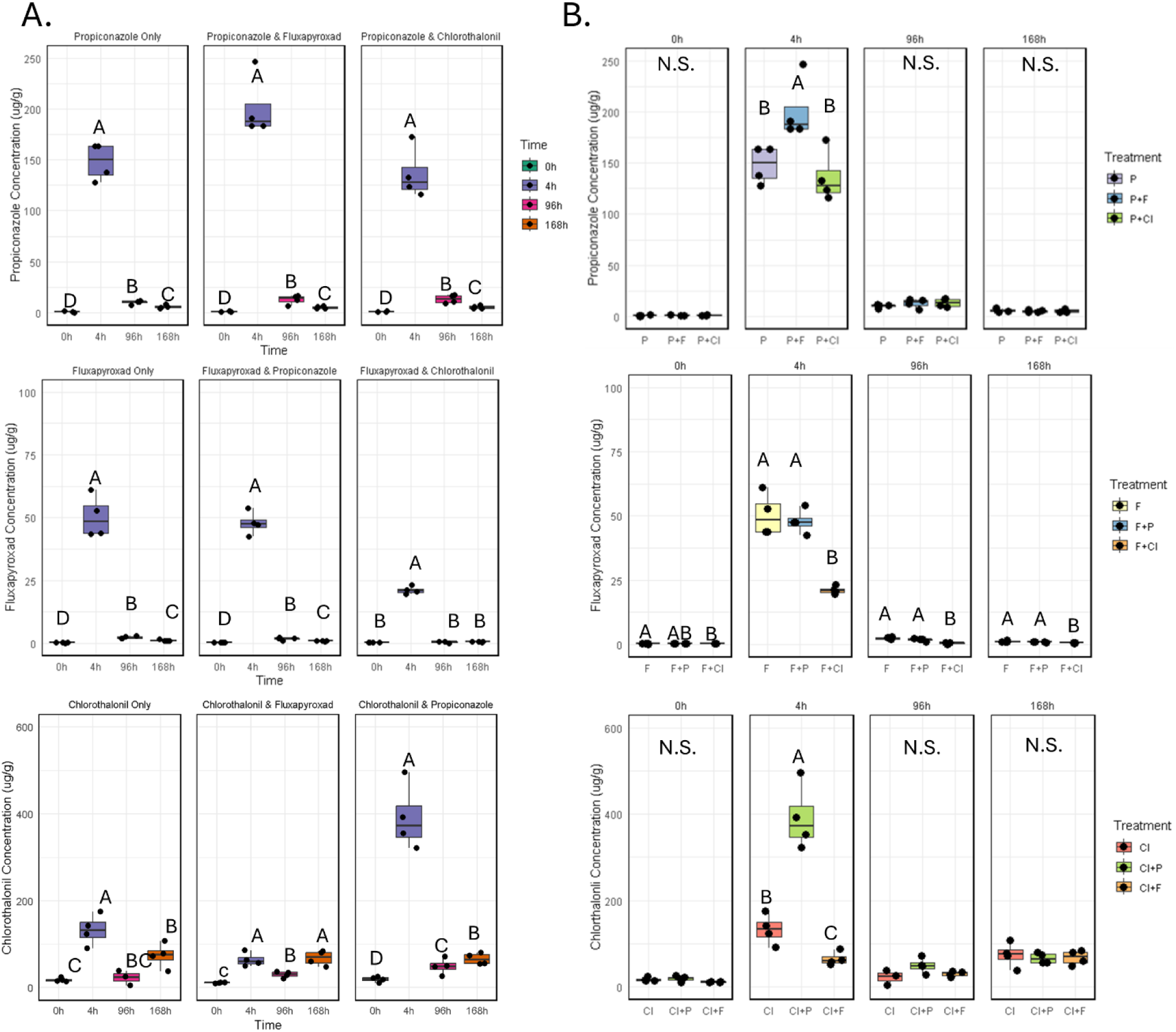
Fungicide active ingredient concentrations vary across time and treatment groups. A. propiconazole, fluxapyroxad, and chlorothalonil concentrations at 0 hours (0h), 4 hours (4h), 96 hours (96h), and 168 hours(168h) post fungicide application faceted by treatment group. B. Propiconazole and Fluxapyroxad concentrations between plots treated with Propiconazole (P), Fluxapyroxad (F), and Chlorothalonil (Cl) as singular treatments or in combination faceted by time. Letters indicate significance groups (p<0.05) within each panel paired (time) or unpaired (treatment) Welch’s T-tests. Points represent the average concentration of each plot from triplicate or duplicate injections. Each boxplot contains 3-4 points based on turf availability.

**Figure 5.**
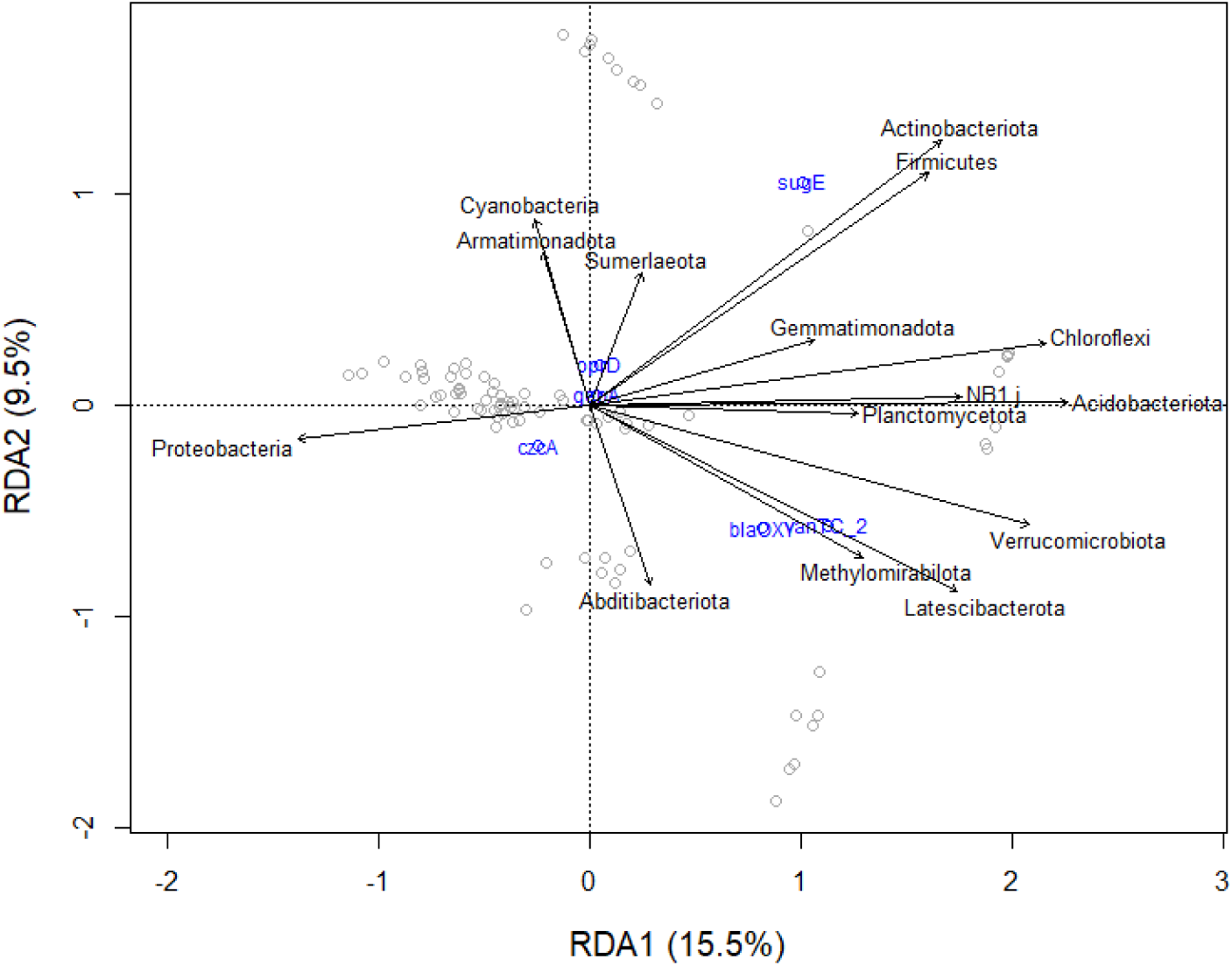
Redundancy Analysis (RDA) biplot for the quantitative correlation of the relative abundance of phyla (exploratory variable) on the log relative abundance of ARGs and MRGs in turf (response variable). Exploratory variables are represented as vectors while response variables are represented as points. Type II scaling was used to represent correlations between exploratory and response variables.

### Fungicide active ingredient concentrations were impacted by time and treatment group

The lack of treatment effect on both the resistome and the bacterial community composition prompted us to measure the concentration of fungicides in our turf samples at each timepoint to ensure that we were seeing differences in active ingredient concentrations. For all fungicides the highest concentration of active ingredients was detected 4 hours post application with average concentrations of 159.5 µg/g, 36.5 µg/g, and 195.7 µg/g for propiconazole, fluxapyroxad, and chlorothalonil respectively. By one-week post application concentrations for propiconazole and fluxapyroxad fell back down to average concentrations of 5.4 µg/g and 1.5 µg/g. Chlorothalonil concentrations at 96 hours averaged 34.6 µg/g and increased at 168 hours to an average concentration of 69.9 µg/g.

Combined treatments did lead to some significant differences in active ingredient concentrations. Propiconazole concentrations at 4 hours were significantly higher when combined with fluxapyroxad compared to the other treatments. However, by 96 hours post application propiconazole concentrations are statistically similar across treatment groups. Comparisons of fluxapyroxad concentrations between treatment groups show significantly lower concentrations when combined with chlorothalonil samples across all time points. Lastly, chlorothalonil concentrations vary significantly between all of the treatment groups at the 4 hour time point. It is unclear if differences between treatment groups are the result of increased variability at the 4 hour timepoint due to increased concentration, matrix effects impacting the instrument’s measurements, or altered degradation rates for treatments applied as mixtures in the field.

### Bacterial community members correlate to ARGs and MRGs in the phyllosphere

The shared temporal trends in the resistome and bacterial community composition data led us to question if specific bacterial community members correlated to ARGs and MRGs. To ascertain relationships between ARGs, MRGs, and bacterial community composition, redundancy analysis (RDA) was performed using phylum relative abundances and log relative abundances of the antibiotic resistance genes, model testing led to the inclusion of phyla *Proteobacteria, Latescibacterota, Gemmatimondota, Methylomirabilota, Chloroflexi*, *Verrucomicrobiota, NB1. J, Acidobacteriota, Plantomycetota, Adidtobacteriota, Cyanobacteria, Armatimonadota, Sumerlaeota, Actinobacteriota,* and *Firmicutes,* within the model as exploratory variables. This model accounts for a total of 32.84% of variation in the log relative abundance of resistance genes within the data set. Permutation tests indicate that *Proteobacteria, Latescibacterota*, *Chloroflexi*, and *Actinobacteria* have significant (p<0.05) impacts on the overall relative abundance of resistance genes independent of model order. These results suggest that bacterial community composition at the phylum level have significant correlations to the relative abundance of resistance genes within the phyllosphere.

Spearman correlations were performed to assess pairwise associations between ARGs, MRGs, and bacterial genera. Results show significant (p <0.05) but moderate (0.4< |ρ| <0.6) correlations between the resistance genes and genera, suggesting that some members within each genus may have stronger correlations with the resistance genes than others. Multiple genera from the phyla *Proteobacteria* and *Actinobacteria* were shown to have significant spearman correlations with *czcA, qepA,* and *oprD. Proteobacteria* and *Actinobacteria* were also found to have significant impacts on resistance gene abundances according to the RDA. Additionally, there are significant pairwise correlations between resistance genes and genera from the phylum *Bacteriadota. Bacteriadota* was not included in the RDA based on model testing, which may have been due to the emphasis on linear relationships in the RDA, reduced significance in the multivariate data set, or multicollinearity with other phyla. We also see that ARGs have significant and mostly strong (0.75< |ρ| <1.0) positive correlations to each other. However, the metal resistance gene (*czcA)* has only moderate (0.4< |ρ| <0.6) positive correlations with ARGs *oprD* and *qepA*. All significant spearman correlations are shown in Figure 6.

**Figure 6.**
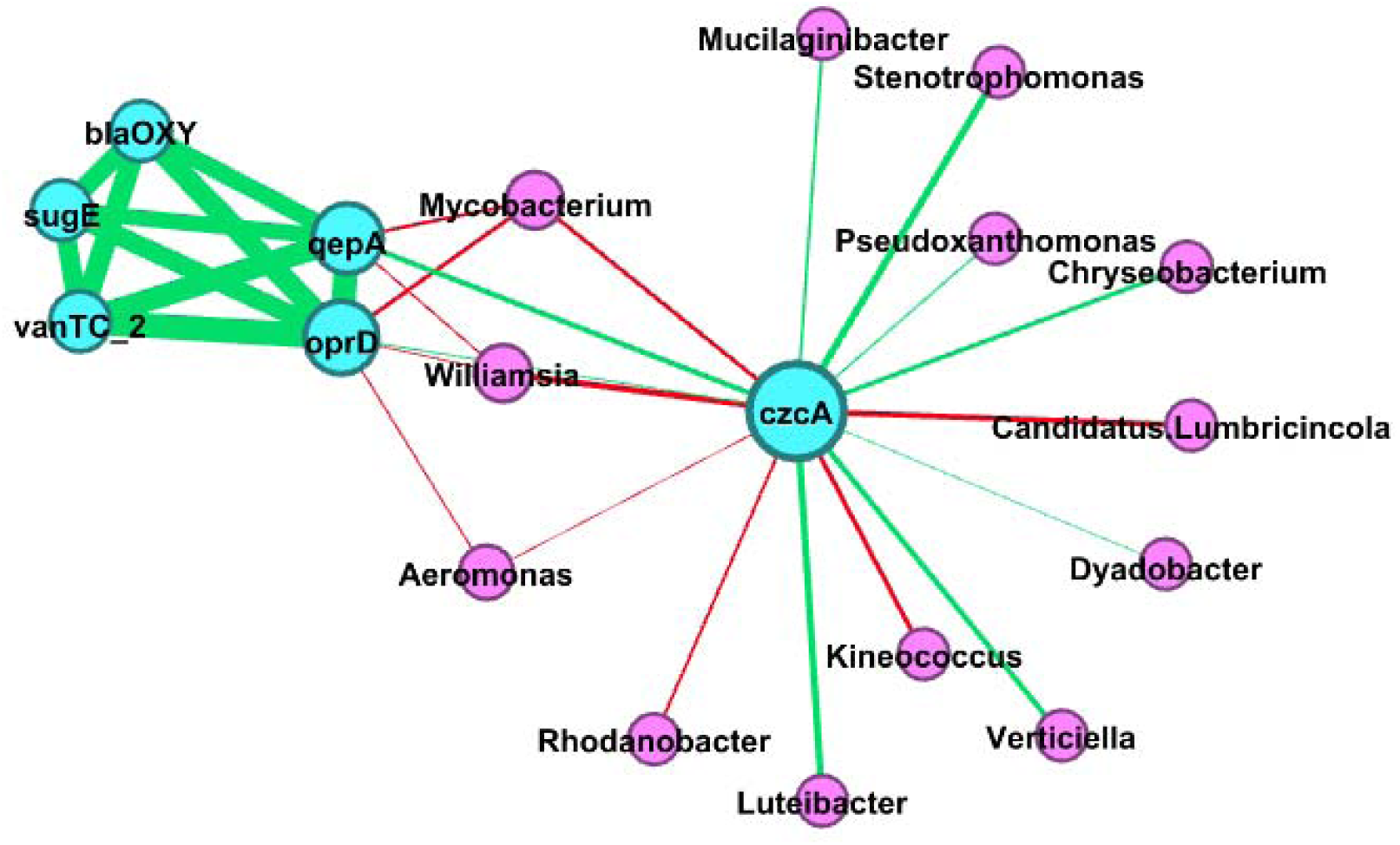
Network representing significant (p < 0.05) pairwise spearmen correlations between ARGs, MRGs, and genera using a Yifan Hu layout. Edges represent correlations that are either negative (red) or positive (green), with the edge weight representing rho. Nodes represent either genus (pink) or MRGs and ARGs (blue), with node weight representing betweenness centrality.

## Discussion

The abundance of ARGs and MRGs was not significantly impacted by treatment but did change with time. The log relative abundance of resistance genes in turf ranged between 10^-2^ and 10^-4^, this range is in line with the relative abundances of resistance genes found in other studies conducted on plants, soil, and water (12, 56–60). The presence of these genes within bacterial communities appears to be relatively low overall within environmental compartments and become more sparse moving up the soil to plant continuum (61, 62). Previous studies have seen shifts in ARG abundance based on treatment with fungicide, but response varies with the ARGs measured and fungicides used. A study conducted on greenhouse soils and mountain soils saw that 60 days after exposure to fungicides, including chlorothalonil and a triazole fungicide, there were significant differences in relative abundance of ARGs compared to non-treated soils, but the effect varied based on the soil type, gene measured, and fungicide applied (12). Additionally, earthworms in chlorothalonil-treated soil were also shown to have an overall increase in abundance of ARGs compared to those in untreated soil 28 days after the addition of fungicide. Worms in the treated soil had significantly increased relative abundance of peptide and beta-lactam resistance genes but significantly reduced the relative abundance of rifamycin, tetracycline and glycopeptide genes in their guts (13). These results show that fungicide exposure can alter ARG abundance in soil systems, but that these shifts in abundance are also dependent on other environmental factors and the genes measured. The lack of treatment effect within our study may show that the phyllosphere resistome is more resilient to short term fungicide application periods than those of other environments, which may be a result of the low bacterial density and metabolic activity within the phyllosphere. Additionally, the overall low relative abundance of resistance genes within the phyllosphere may limit our ability to detect changes driven acute exposure to fungicides. The change in abundance of ARGs over time, but not in response to treatment, suggests that shifts in the bacterial community composition or other environmental factors may have a larger impact on the resistome than fungicide exposure within the timeframe of our study.

Applications of fungicides failed to significantly alter the phyllosphere bacterial community. Previous studies have similarly found no significant changes in bacterial community composition in response to short term fungicide exposure, however, the phyllosphere has been understudied and turf overlooked. A lack in changes to community composition in response to azole fungicides, like propiconazole, have previously been demonstrated in the phyllosphere of vegetables (63, 64). The impact of short-term chlorothalonil and fluxapyroxad application on the composition of phyllosphere bacterial communities were unknown but results similar to our findings were shown in soil. Turf soils treated with chlorothalonil were shown to have no significant difference in bacterial community composition when compared to untreated soils two weeks post application (16). Additionally, fluxapyroxad has previously been shown to have no impact on the bacterial communities of soil at lowbush blueberry agricultural sites after a 24-day fungicide trial (15). However, to our knowledge, studies of combined fungicide treatments have not examined impacts to the bacterial community in any plant niche. The lack of change to the bacterial community in these field experiments suggests that bacteria are resilient to fungicide treatment after short exposure periods, but it’s unclear whether this stability persists over long time periods.

Fungicidal effects on bacterial community composition may change over prolonged application periods. For example, Li, et al. (2023) periodically applied propiconazole and chlorothalonil as singular treatments and in combination with each other over a four-month period on boxwood shoots and the impact of fungicides on the epiphytic bacterial community varied over the application period (65). Through their early summer sampling on days 44 through 58, no significant difference in the beta diversity of epiphytic bacterial communities among their treatment groups was observed. However, there were significant differences based on the time point over the course of the two weeks when sampling occurred. These early application results largely reflected our findings where treatment did not have an impact, but sampling time did. Li, et al. (2023) then found in the early fall sampling, after fungicides had been applied periodically over 135 days, treatment had a significant impact on the beta diversity of boxwood shoot epiphytes while the sampling time was no longer significant. Changes to beta diversity after long term management differences more closely reflected the findings of other studies performed by our lab, which found significant differences in phyllosphere bacterial community composition after long-term management intensities differences in turf (66, 67). These results suggest that treatment impacts on the bacterial community composition of the phyllosphere may emerge over extended application periods and that sampling timepoint may become less significant.

Fungicide active ingredient concentrations decreased 64.3 - 96.6% from the 4 hour to 168 hour timepoints. Previously, propiconazole and chlorothalonil have been found to have half-lives ranging 12 to 15 and 4.0 to 4.3 days respectively in creeping bent grass (68, 69). The half-life of fluxapyroxad in turf is unknown, but fluxapyroxad has been shown to have a half-life of 2.6 and 12.2 days on cabbage and scallions respectively (70). In our study, chlorothalonil concentrations increased from the 96 hour timepoint to the 168 hour timepoint, this may be due to relying on in-source generation of 4-hydroxyl-chlorothalonil as opposed to native chlorothalonil in our LC-MS/MS analyses. Potter et al. (2001) has shown that 4-hydroxyl-chlorothalonil accumulates in soil over five days post foliar application to peanuts, though it is unclear if 4-hydroxyl-chlorothalonil is also capable of accumulating in plant tissues (71). While the chemistry of chlorothalonil in-source transformation to 4-hydroxyl-chlorothalonil has been documented (72), the mechanism is not well understood and consequently the conditions that yield predictable in-source transformation remain elusive. Considerable work remains to fully characterize the robustness of chlorothalonil measurements in plant tissue by LC-MS/MS. However, we did see significant changes in all of the active ingredient concentrations over time, suggesting that bacteria were exposed to different active ingredient concentrations despite the lack of treatment effect on community composition and resistance gene abundances.

The agreement in temporal trends between resistance gene abundance and beta diversity data prompted us to question whether there are correlations between ARGs and bacterial community members. Significant correlations between phylum *Proteobacteria, Latescibacterota*, *Chloroflexi*, and *Actinobacteria* and resistance genes found via redundancy analysis suggest that these community members correlate to the overall abundance of the five measured ARGs and one MRG in significant and predictable ways. Redundancy analysis is limited due to linearity assumptions, potentially explaining why we see different relationships when running pairwise spearman correlations. The resistance genes we measured had significant spearman correlations to thirteen different genera across five different phyla. These results are consistent with previous studies that have found that bacterial community structure correlates with the abundance of ARGs. Specifically, *Actinobacteria*, *Firmicutes,* and *Proteobacteria* have been found to correlate to the presence of ARGs in soil, earthworm guts, wastewater treatment plants, and tree leaves (13, 73–77). In our study, *czcA* was positively correlated with *Dyandobacter*, *Verticiella, Chryseobacterium, Muclilaginibacter,* as well as the opportunistic human pathogens *Psudoxanthomonas, Stenotrophomona*s, and *Luteibacter*. Positive correlations with these genera suggest that they may act as hosts for *czcA* within this environment. Additionally, ARGs had strong (0.75< |ρ| <1.0) positive correlations with each other as has been previously seen in soil and wastewater, with other ARGs (7, 39, 78). We also saw ARGs *oprD* and *qepA* correlate positively with the MRG *czcA*. Previously, czcA has been reported to correlate with the ARG *mphB* in wastewater (39). The positive correlation between resistance genes may indicate co-selection of these genes or their presence on shared mobile genetic elements.

Correlations between taxa, ARGs, and MRGs suggest that bacterial community composition has a significant relationship with the resistome of the phylloshere, similarly to other environmental niches. It is unclear if the lack of treatment effect within this study is due to a unique resilience of the phyllopshere resistome, unique attributes of the turfgrass system, or due to the short exposure period and genes measured in this study. Future studies incorporating longer fungicide application periods and more ARG targets would help to clarify if treatment effects on ARG abundance emerge in the phyllosphere under different experimental conditions.

## Acknowledgements

This material is based upon work supported by USDA Hatch Project WIS303044. The funders had no role in study design, data collection, and interpretation or the decision to submit the work for publication.

We thank Nathan Aviles and Nicholas Keuler from the Statistical Consulting Group at the University of Wisconsin–Madison for assistance in the analysis of the ARG abundance data. Additionally, we would like to thank Finn Kuusisto for assistance with the network analysis.

## Notes

### Competing Interest Statement

The authors have declared no competing interest.

### Summary of Updates

Based on the responses to reviewers, the manuscript underwent substantial revision to improve clarity, methodological transparency, and interpretation of results. We strengthened the Introduction by expanding the rationale for studying resistance genes in the phyllosphere, clarifying how plant-associated microbial communities differ from soil environments, and more explicitly stating the study hypotheses and experimental approach. Terminology was standardized throughout (e.g., replacing 16s with 16S rRNA), and descriptions of fungicide modes of action, sampling procedures, and transport/storage methods were clarified. Methodological details were significantly expanded in response to reviewer concerns. We provided additional information on the high-throughput qPCR approach, including use of real-time qPCR chemistry, sample preparation, primer validation, Ct filtering criteria, and calculation of relative abundance. References and descriptions for primer design, bioinformatics workflows, statistical modeling, and software versions were added. We also clarified the distinction between technical and biological replication and improved reporting of statistical results by including treatment-associated p-values. To improve presentation of results, a figure illustrating temporal changes in resistance gene abundance was moved from the supplemental material into the main manuscript, while a figure considered uninformative was removed. Additional analyses evaluating sequence coverage were conducted, and Goods coverage metrics were added to demonstrate adequate sampling depth for microbial community analyses. The Discussion was revised to better acknowledge study limitations and avoid overinterpretation. We tempered language regarding correlations between bacterial taxa and resistance genes, emphasized that correlation does not imply causation, and expanded discussion of limitations associated with short-term sampling, single fungicide applications, and the relatively low abundance of resistance genes in the phyllosphere. We also highlighted the need for longer-term studies evaluating repeated fungicide exposure. Finally, the manuscript title was revised from Time, not fungicide treatment, influences the resistome of the turf phyllosphere to Time influences the resistome of the turf phyllosphere to better reflect the scope of the data and avoid overstating conclusions regarding fungicide effects. Collectively, these revisions improve the manuscript rigor, transparency, and balance while maintaining the central finding that temporal shifts in microbial community structure were more strongly associated with resistome dynamics than acute fungicide exposure.

