## Supplementary material for "Temporal Dynamics Outweigh Fungicide Treatment in Shaping the Turfgrass Phyllosphere Resistome": Table S1-S2 and Fig. S1-S3

### Supplementary Materials

Supplementary Table 1. LC-MS/MS Gradient Method for Chlorothalonil Quantitation

| Time (min) | A (%) | B (%) |
|---|---|---|
| **0** | 90.00 | 10.00 |
| **0.5** | 90.00 | 10.00 |
| **5.00** | 0.00 | 100.00 |
| **6.00** | 0.00 | 100.00 |
| **6.10** | 90.00 | 10.00 |
| **8.00** | 90.00 | 10.00 |

Supplementary Table 2. Chlorothalonil Source Parameters

| Parameter | Negative Polarity Value |
|---|---|
| **Gas Temperature (°C)** | 230 |
| **Gas Flow (L/min)** | 20 |
| **Nebulizer (psi)** | 35 |
| **Capillary Voltage (V)** | 1950 |
| **Sheath Gas Temperature (°C)** | 370 |
| **Sheath Gas Flow (L/min)** | 9.5 |
| **Nozzle Voltage (V)** | 1700 |

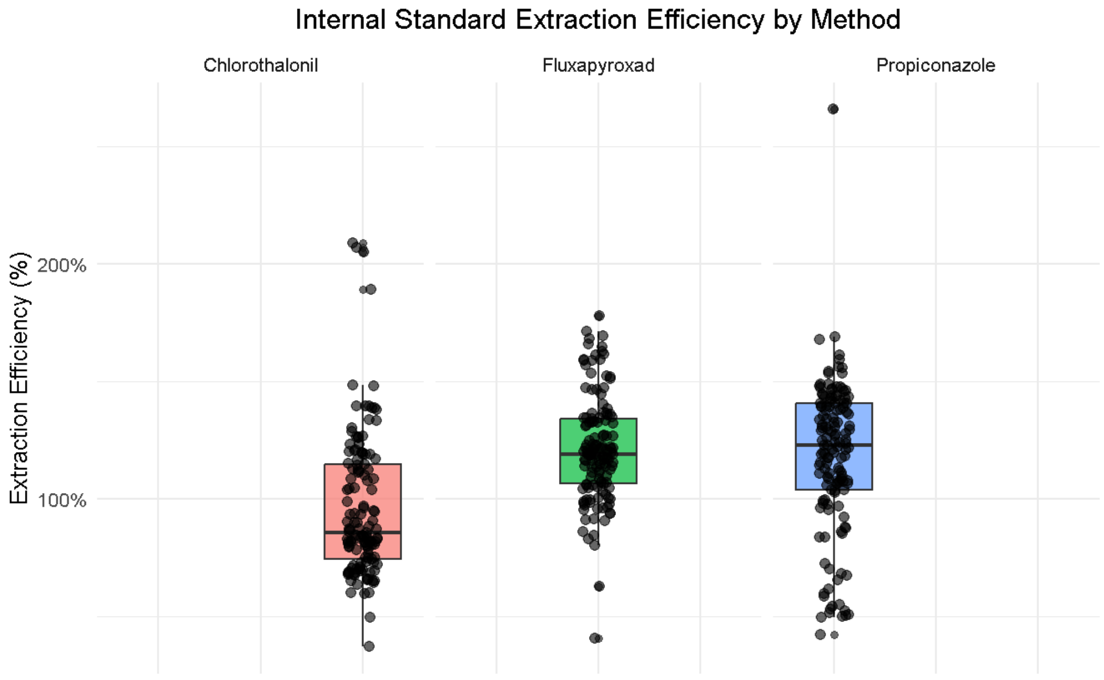


Supplementary Figure 1. Internal standard extraction efficiency by LC-MS/MS method.

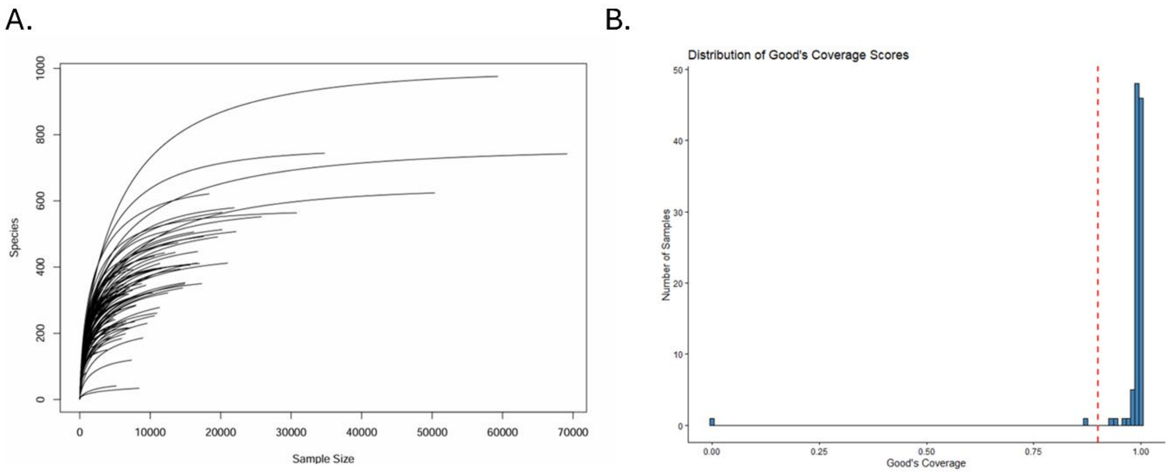


Supplementary Figure 32. Quality Filtered 16s Coverage Results. A Rarefaction Curve at ASV Level. B.  Distribution of Good's Coverage Scores Prior to Normalization. Red dashed line indicates 90% coverage.

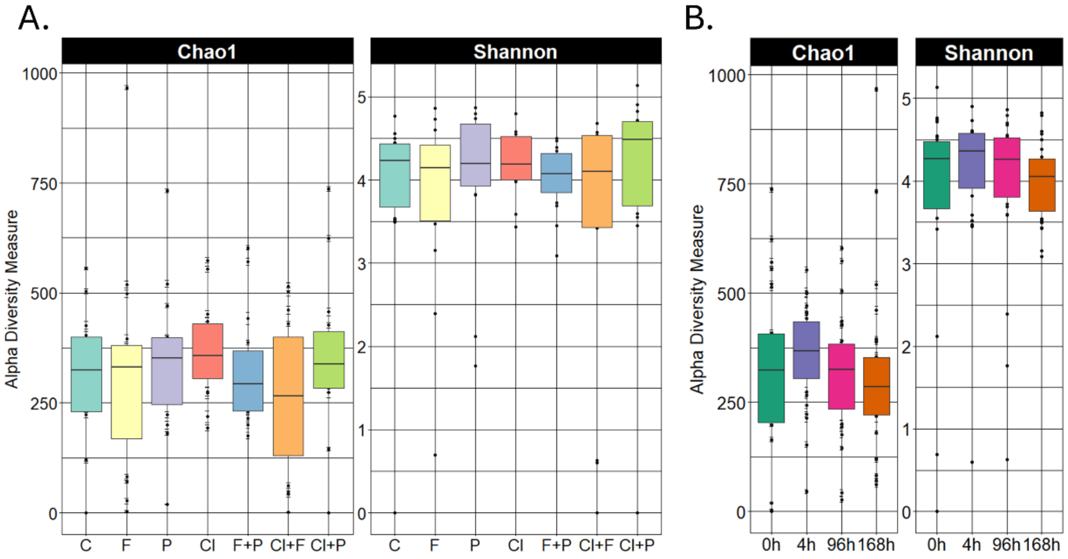


Supplementry Figure 3. Alpha diversity represented by Chao1 and Shannon Indeces. A. Alpha diversity of samples at 0 hours (0h), 4 hours (4h), 96 hours (96h), and one week (168h) post fungicide application, samples are combined across treatment groups. B. Alpha diversity of samples in the control plot (C) and plots treated with fungicides containing Propiconazole (P), Fluxapyroxad (F), and Chlorothalonil (Cl) as singular treatments or in combination, samples are combined across time points.
